# Sex specific knee joint soft tissue mineralization with Fibrillin-1 mutation in male Tight Skin mice

**DOI:** 10.1101/2024.08.28.610095

**Authors:** Craig Keenan, Xi Wang, Tayfun Dikmen, Yan Wen, Lorenzo Ramos-Mucci, Emily Shorter, David Abraham, George Bou-Gharios, Blandine Poulet

## Abstract

Articular soft tissue mineralization and ossification are clear pathological signs of osteoarthritis joints. However their molecular and cellular aetiologies remain largely unknown. TGF-ß family members are known contributors to both pathological ossification and osteoarthritis development. In this study, we used a Fibrillin-1 mutant mouse, the Tight Skin mouse (TSK), to define the detrimental effects of abnormal Fbn1 in TSK mice and known high TGF-ß activity in joint pathology such as articular soft tissue mineralization and ossification.

Knee joints of male and Female TSK and Wild-type (WT) littermates were analysed my micro-CT imaging and histology for articular soft tissue pathologies, as well as OA severity. Both aged (10, 26, 35 and 52wks) and following in vivo non-invasive repetitive joint overloading were used.

We find that male TSK mice develop spontaneous soft tissue ossification from 26wks of age, followed by increased osteoarthritis at 1 year-old. In addition, knee joint overloading induced ligament and meniscal mineralisation and ossification in both WT and TSK male mice, but were significantly more severe in TSK knees, including ossification of the patella ligament and synovial lining. In contrast, female TSK knees did not develop more severe soft tissue mineralisation compared to littermate WT mice in neither aged nor overloaded knees.

We conclude that Fbn1 mutation, and possibly overactive TGF-ß activity in TSK mice, induce articular soft tissue ossification and osteoarthritis in a sex-specific manner. Further studies are needed to confirm the specific signalling involved and the relative protection from female mice from such pathologies.

## Introduction

Mineralization is an important biological process which is responsible for the development of tissues such as bone, cartilage and teeth, as well as their regeneration. Yet, this biological process can also occur pathologically in extra-skeletal tissues (Tsolaki and Bertazzo, 2019). Pathological soft tissue mineralization is a major process involved in heterotopic ossification or as a consequence of diseases such as atherosclerosis (Jeziorska *et al*., 1998), ankylosing spondylitis (Jo *et al*., 2022), tendinopathy (Richards *et al*., 2008) or in joints during osteoarthritis (OA)(Ramos-Mucci *et al*., 2020; Schulze-Tanzil, 2019). Pathological soft tissue mineralization often occurs due to disturbance in the physiological tissue repair process, and most commonly occurs following a traumatic injury (Meyers *et al*., 2019). As a response to trauma, a series of signalling events occur in the injury site which trigger migration of inflammatory cells and mesenchymal cells to facilitate tissue repair. However, excessive TGF-ß activity may disrupt this physiological process and cause activation of osteogenic or osteochondral programmes, and trigger mineralization and extra-skeletal bone formation (Wang *et al*., 2018).

OA is a complex degenerative disease that affects the whole joint and is characterized, but not limited, by cartilage defects, subchondral bone remodelling, osteophyte formation, and ligament degeneration (Loeser *et al*., 2012). Although degenerative articular cartilage is considered the most common outcome of OA, mineralization and pathological endochondral ossification of the soft tissues in the diseased joint, especially in the anterior cruciate ligament (ACL), synovium, and menisci also contribute to OA severity and pain (Ramos-Mucci *et al*., 2020; Schulze-Tanzil, 2019). Understanding molecular and cellular mechanisms of articular soft tissue ossification might represent a novel therapeutic target to slow OA progression.

Fibrillin1 (Fbn1) is a 350 kDa protein is a major structural component of the extracellular matrix (ECM) of connective tissues with high expression in elastic fibres (Godwin *et al*., 2019; Sakai *et al*., 1986). The structure and binding affinities to cytokines, other ECM proteins and cell surface receptors strongly support multiple roles in tissue integrity and mechanical properties and in cell signalling (Halper and Kjaer, 2014). In particular, Fbn1 microfibrils provide a reservoir for latent TGF-ß in the pericellular matrix (Asano *et al*., 2022). This role of Fbn1 is exemplified by the high levels of active TGF-ß in Marfan Syndrome tissues and in murine pre-clinical models with FBN1 mutations, such as the Tight Skin mouse (TSK) (Lemaire *et al*., 2006; Saito *et al*., 1999). The latter harbours an autosomal dominant mutation associated with a duplication of FBN1 gene (Siracusa *et al*., 1996), and in addition to its characteristic thick and stiff skin, TSK mice also exhibit Marfan-like pathologies such as bone overgrowth, cardiac hypertrophy, emphysema-like lungs and kyphosis (Barisic-Dujmovic *et al*., 2007; Green *et al*., 1976). The close relationship of Fbn1 with TGF-ß superfamily proteins, ECM formation and structure is thought to be a major contributor to the musculoskeletal phenotype of TSK mice.

The general aim of our study is to define the detrimental effects of abnormal Fbn1 in TSK mice and known high TGF-ß activity in joint pathology such as articular soft tissue mineralization and ossification. Herein we show that knee joints of TSK mice develop sex-specific mineralization and ossification in ligaments and synovial tissues, exacerbated by in vivo joint overloading.

## Materials and Methods

### Animals

Male and female Tight Skin (TSK) mice imported from The Jackson Laboratory (Bar Harbor, ME) and maintained as a colony at the University of Liverpool. Homozygote TSK mice are not viable, and therefore all mice referred to TSK are all heterozygotes for the mutated Fbn1 gene. To avoid potential breeding issues, breeding pairs consisted of a male TSK and a WT female, ideally from the same litter. All mice were kept in polypropylene cages of 2–5 mice, subjected to 12-hour light/dark cycles at 21±2°C, with free access to food (pelleted RM1; SDS diets) and water. All procedures complied with Animals (Scientific Procedures) Act 1986, local ethics committee and under appropriate Home Office project licences. Both males and females were left to age to 10 and 26 weeks of age, and to 35 (females due to COVID restrictions) or 60 weeks of age (males). Animal numbers for each group are included in Table 1. Mice were killed by increasing amounts of CO_2_, and knees collected.

**Table 1:** Animal numbers used.

| **Males** | **10wks** | **26wks** | **60wks** | **Loading** |
| --- | --- | --- | --- | --- |
| WT | 10 | 11 | 4 | 13 |
| TSK | 7 | 14 | 4 | 17 |
| **Females** | **10wks** | **26wks** | **35wks** | **loading** |
| WT | 9 | 11 | 18 | 13 |
| TSK | 8 | 12 | 14 | 14 |

### Joint overloading

The right knee of male and female mice (WT and TSK littermates) were mechanically loaded repetitively, as described before (Poulet *et al*., 2011). Briefly, mice were kept anaesthetised throughout each loading episode (∼7 minutes), which consisted of 40 cycles of 9N loads (peak load time: 0.05sec; rise and fall time: 0.025sec), with a 2N holding load in between of 9.9sec, using the ElectroForce 3100 system (TA Instruments, USA). Six loading episodes were performed over 2 weeks, and knee joints were collected 6 weeks after the last loading episode. We showed that this time point is sufficient to induce soft tissue mineralization in the ligaments and meniscus (Ramos-Mucci *et al*., 2022). Mouse weights were not different with genotype (mean ± standard deviation: males WT: 39.65g±4.45; Male TSK: 39.9g±4.41; Female WT: 33.45g±4; Female TSK: 30.42±5.77). Animal numbers can be found in Table 1.

### Micro-Computed Tomography (µCT)

Following fresh tissue collection, knees were fixed in 10% neutral buffered formalin for 24-48h, then transferred to 70% ethanol. We used a protocol used previously (Ramos-Mucci *et al*., 2020). Knees were scanned at a resolution of 4.5 µm using a 0.25 mm aluminium filter, with a rotation step of 0.6° (Skyscan 1272; Bruker micro-CT, Belgium). Image reconstruction was performed using NRecon software (Bruker micro-CT, Belgium), followed by manual selection of regions of interest for the joint space mineralization volume using CTAn (Bruker micro-CT, Belgium), as described (Ramos-Mucci *et al*., 2020), which included meniscal tissue and any abnormal mineralized tissues that was not tibial or femoral bone. Mineralized tissue volume was calculated using the 3D algorithm included in the CTAn software (measured as Bone Volume). Statistical analysis of mineralized volume comparing the different groups was performed using one-way ANOVA with Tukey’s post hoc test; to test the significance of the effect of joint loading, paired t-test were performed for each genotype between the Left non-loaded Contralateral knee and the Right Loaded knee. Three-dimensional models of the menisci were created using CTVox from the region of interest selected for mineralized tissue volume analysis (Bruker micro-CT, Belgium).

### Histology

Knee joints were decalcified in 10% formic acid (Sigma-Aldrich, UK) for 10 days. Sample were then given a processing number independent of their genotype and processed for wax embedding, and serial sections cut at 6-µm-thick either the coronal (ageing) or sagittal plane (loaded samples) across the entire joint. Sections at 120 µm intervals were stained with Toluidine Blue/Fast Green (0.04% in 0.1 M sodium acetate buffer, pH 4.0). Pathological assessment of all joints was performed, focusing on the soft tissues within the joint (including ligaments and synovial lining). In addition, cartilage lesion severity was graded in ageing samples using the Osteoarthritis Research Society International (OARSI) histopathology initiative scoring method (Glasson *et al*., 2010). Grading each of the four compartments of the tibiofemoral joint (lateral and medial tibia and femur) throughout the entire joint allowed for the determination of average (mean) lesion grades for each condyle, and added to make a summed mean score for each joint (Poulet *et al*., 2011). Statistical analysis was performed using one-way ANOVA with Tukey’s multiple comparison test. Data are presented as box plots of interquartile range, median, minimum and maximum, showing all individuals.

## Results

### Male TSK mouse knees developed spontaneous soft tissue ossification that precedes osteoarthritis development

Male TSK mice showed significantly increased soft tissue mineralisation from 26 weeks of age, measured by µCT as a volume of intra-articular joint space (Fig.1), with volumes increasing from 0.239×10^9^ (±0.013 sem) µm^3^ in WT male mice at 26wks to 0.801×10^9^ (±0.218 sem) µm^3^ in TSK mice (p=0.0502). Significant mineralised tissue mass was formed in the anterior part of the knee joint, as well as some nodules around the meniscus (Fig.1B). Toluidine blue staining of histological sections supports cartilage and even bone formation in the patella ligament (Fig.1C-E). In the severely affected joint, the meniscal attachments to the femur and tibia, as well as the meniscal ligaments and synovial tissue, formed proteoglycan-rich ECM with areas of clear ossification (Fig.1F-G)

**Figure 1:**
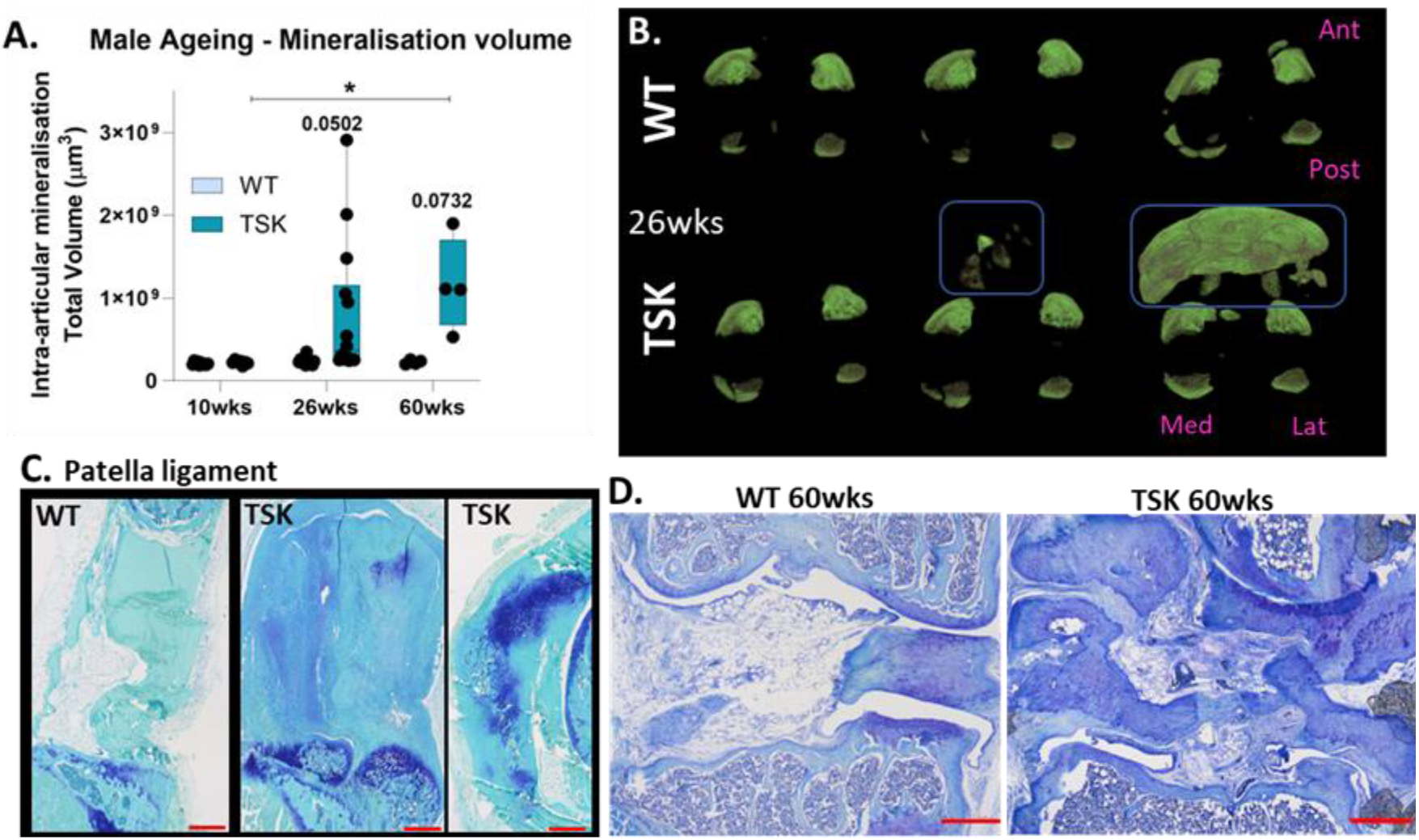
Spontaneous soft tissue mineralisation and ossification in Male TSK mice. **A**. Total volume of intra-articular mineralisation measured by µCT image analysis at 10wks, 26wks and 60wks of age; * for p<0.05. **B**. 3D volumetric representation of intra-articular mineralisation in 26wk-old WT and TSK male mice, proximal view of the smallest, median and highest volumes in each group. Blue circle highlight the severe anterior mineralisation seen in TSK mice. **C-D**. Coronal toluidine blue stained knee joints from representative samples from C) 26wk-old patella ligaments in WT and TSK mice showing signs of cartilage and bone formation, D) and from 60wk-old WT and TSK showing severe cartilage and bone formation around the meniscus, in the ligaments and synovial tissues (red scale bar represents 50µm).

The OARSI scoring showed that male TSK mice developed spontaneous osteoarthritis compared to WT controls (Fig.2), from a summed OARSI score across all 4 condyles of the knee in WT at 60wks of age of 2.0 (±0.196) to a score of 6.9 (±2.13; p=0.0015). While detrimental pathological changes were observed in male TSK mice at 26 weeks, no cartilage degeneration was seen at this age.

**Figure 2:**
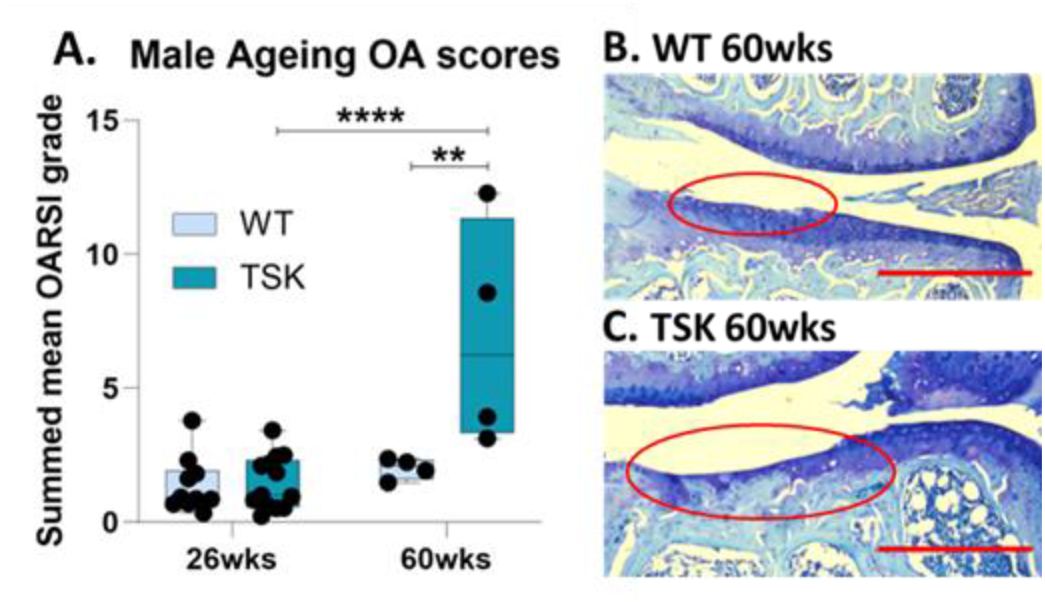
Spontaneous OA development in male TSK mice. **A**. Summed mean OARSI cartilage degeneration scores across the whole joint; **** for p<0.0001. **B-C**. Representative images of Toluidine Blue stained coronal sections showing articular cartilage degradation (red circle) in the tibia of 60wk-old WT (B) and TSK (C) knee joints. (red scale bar represents 50µm).

### Male TSK mice developed more severe mechanically-induced articular soft tissue ossification

Both WT and TSK mice demonstrated increased intraarticular mineralisation after repetitive mechanical joint loading, which was quantified and visualised with µCT (Fig.3A-B). Loading-induced nodule were seen around the meniscus (red arrows Fig.3A) in both WT and TSK, with more severe volumes observed in TSK mice, as well as additional mineralised nodules across the joint, including in the anterior compartment of the patella ligament. In WT mice, mineralised volume increased by 0.052 µm^3^ from 0.233 (±0.006) µm^3^ in non-loaded left legs to 0.286 (±0.008) µm^3^ (p<0.0001) in contra-lateral right loaded knees. In contrast, TSK mice showed an increase of 0.389 µm^3^ from 0.282 (±0.010) µm^3^ in non-loaded left legs to 0.672 (±0.112) µm^3^ (p=0.002) in contra-lateral right loaded knees; loaded right legs showed a significant increase in TSK knees with a p=0.0002. Histological toluidine blue staining also confirmed the severity of mineralisation in TSK mice with clear patellar ligament ossification, osteophyte formation and disorganisation of the enthesis of the patella ligament into the tibia, as well as chondrogenesis in the synovium (Fig.3C-E). These suggest a potential mechanical aetiology of these soft tissue ossification processes in TSK mice.

**Figure 3:**
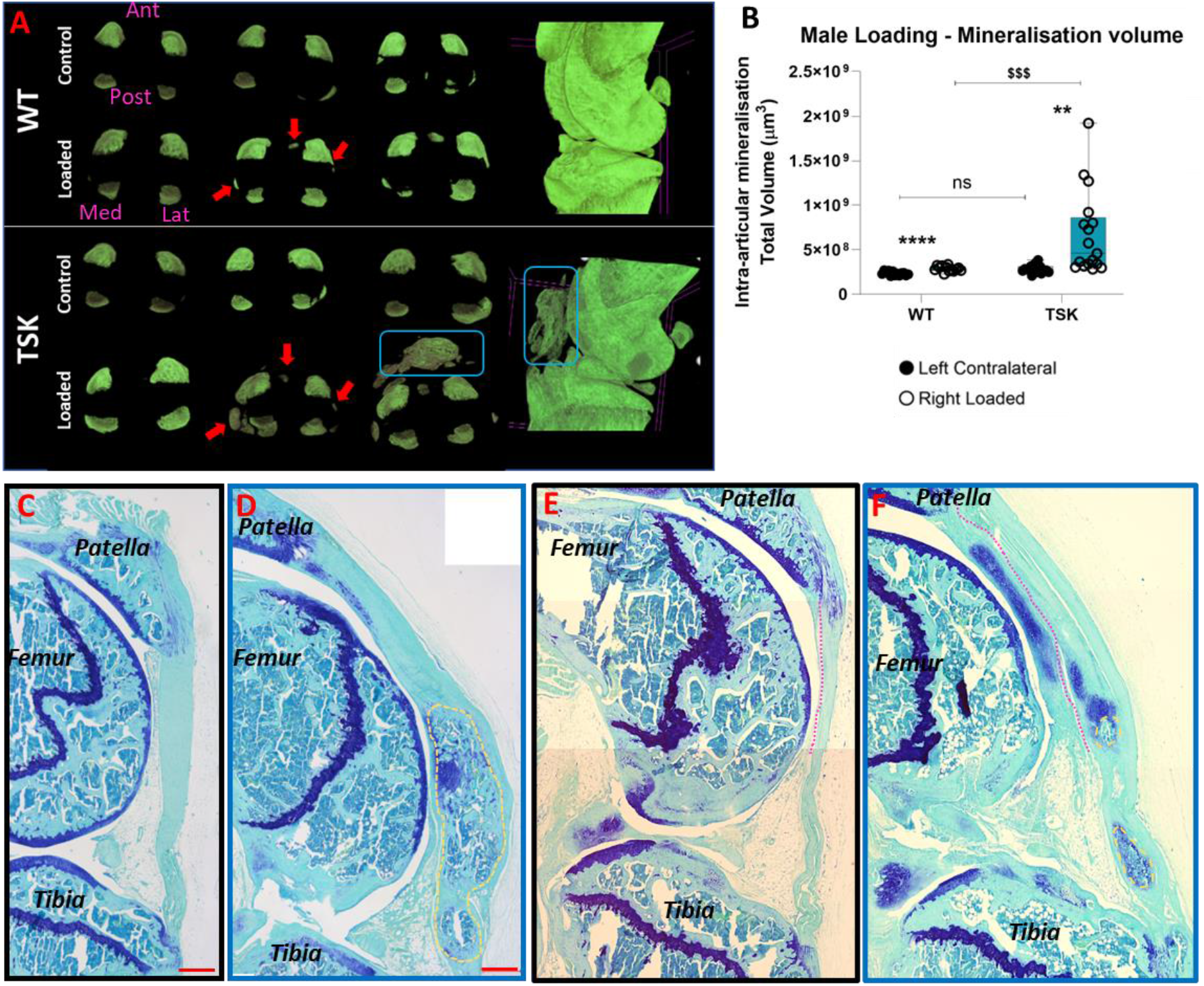
Loading-induced intra-articular soft tissue mineralisation and ossification in male WT and TSK knees. **A**. 3D image representation of the volume of mineralised tissues visualised by uCT in WT (top) Control non-loaded and Loaded knees versus TSK (bottom) control non-loaded and loaded knees. 3 examples with the smallest volume, median and highest volumes are represented for each group. Red arrrows show location of typical load-induced mineralisation nodule formation; blue square highlights the severe anterior mineralisation seen in TSK joints. Right images represent 3D images of the whole joint. **B**. Volume of intra-articular mineralisation in male WT and TSK knee joint in response to joint loading protocols; statistical significance: Left versus Right (paired t-test: ** for p<0.01; **** for p<0.0001); ANOVA with post-hoc test between all groups (&&& for p<0.001). **C-F**. Histological sagittal sections of knee stained with Toluidine Blue following joint loading in WT (C, E) and TSK (D,F); yellow circles delineate ossified regions of the patella ligament (D,F); pink line separates synovial lining from patella ligament at the femoral end (E,F).

### Female TSK mice are not different to their WT littermates

Knee joint mineralisation volumes were similarly measured by µCT in female WT and TSK mice with ageing (up to 35wks of age) and in response to joint loading, as performed in male mice. In contrast to male mice, no significant differences with genotype were measured (Fig.4). Both WT and TSK female mice showed a significant increase in intra-articular mineralisation volume in response to joint loading, as seen in male mice, with no effects of genotype seen. In WT mice, mineralised volume increased by 0.238 µm^3^ from 0.208 (±0.007) µm^3^ in non-loaded left legs to 0.447 (±0.072) µm^3^ (p=0.005) in contra-lateral right loaded knees. In contrast, TSK mice showed an increase of 0.392 µm^3^ from 0.228 (±0.004) µm^3^ in non-loaded left legs to 0.620 (±0.080) µm^3^ (p=0.0004) in contra-lateral right loaded knees.

**Figure 4:**
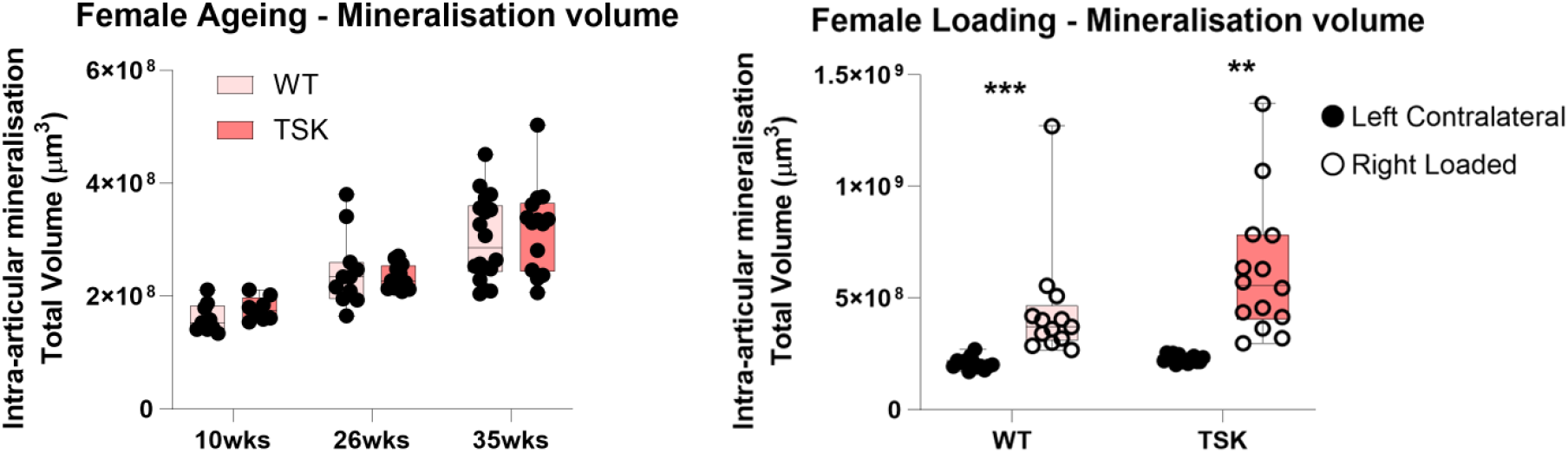
Female TSK do not show increased soft tissue mineralisation compared to WT female mice. A-B. Volume of intra-articular mineralisation in female WT and TSK knee joint at 10, 26 and 35wks of age (A) and in response to joint loading protocols (B); statistical significance: Left versus Right (paired t-test: ** for p<0.01; *** for p<0.001).

## Discussion

FBN1 mutation in TSK mice, linked to increased active TGF-ß, resulted in spontaneous and mechanically-induced articular soft tissue ossification, which precede increased OA development. Interestingly, we show these effects are largely restricted to male mice. This data suggest that abnormal FBN1, such as increased degradation, may contribute to OA articular soft tissue pathologies independently of articular cartilage degeneration. Understanding the sex dimorphism in these responses may also give us clues into sex-specific mechanisms of joint degeneration with important implications for therapeutic targeting of OA progression.

High TGF-ß activity have been linked to heterotopic ossification (Wang *et al*., 2018; Yan *et al*., 2017), which we show to develop in our male TSK knee joints. Fbn1 acts as a regulator for TGF-ß activity with the binding of inactive TGF-ß precursors in latent complexes in close proximity to the cell surface within the pericellular environment. Hence, alterations in Fbn1, as seen in TSK mice and in Marfan Syndrome tissues, result in abnormal TGF-ß activation and is linked to excessive bone growth (Barrett and Topol, 2013). Although the exact mechanism of pathological soft tissue mineralization is still not fully understood, the TGF-ß signalling and increased activity are often considered as one of the most important factors contributing it (Toom *et al*., 2007; Yu *et al*., 2021). Toom et al. (2007) previously reported the presence of TGF-ß superfamily proteins in heterotopic ossification zones, and pointed out the higher bone-forming activity, especially with the involvement of TGF-ß1-3 and BMP-2 in these zones. Wang et al (2018) used transgenic mouse approaches and inhibitory antibody treatment to show the importance of TGF-ß in heterotopic ossification in mouse skeletal tissues including tendons.

In healthy joints, TGF-ß superfamily members control the balance between ECM synthesis and degradation (Blaney Davidson *et al*., 2007a; Finnson *et al*., 2012; Varela-Eirin *et al*., 2018; Zhang *et al*., 2012), are known to promote cartilage formation and repair (Wu *et al*., 2024; Yoo *et al*., 2022; Zhang *et al*., 2015a). Increased OA development in our TSK mice, however, support a detrimental effect of long-term over-active TGF-ß in joints, and suggest therapies aimed at increasing TGF-ß to improve cartilage repair are not suitable for extended periods of time. During OA development, imbalances in growth factor levels have been reported; indeed, elevated TGF-ß levels is linked to increased ECM turnover, osteophyte formation and synovial fibrosis, and therefore promotes disease development (Blaney Davidson *et al*., 2007b; Blaney Davidson *et al*., 2006; Shlopov *et al*., 2001). In addition, it has been shown that the canonical receptors for TGF-ß shifts from an anabolic signalling (via ALK5 – SMAD2/3) to a catabolic pathway (via ALK1 – SMAD1/5/8) (Blaney Davidson *et al*., 2009), thereby increasing cartilage degradation. Understanding the fine tuning of TGF-ß activities, maybe via Fbn1 levels and localisation, may help us develop better therapeutic strategies to prevent or slow OA development. However, it is important to note that the role of TGF-ß activity in the described phenotype in TSK mice has not been assessed in this study.

To further investigate the susceptibility of TSK joints to soft tissue ossification, we used the non-invasive knee joint overloading model, in which repetitive knee compression over 2 weeks leads to increased meniscal and ligament endochondral ossification (Ramos-Mucci *et al*., 2022). FBN1 mutation in TSK mice resulted in enhanced articular soft tissue ossification, confirming increased susceptibility in both spontaneous and mechanical pathological responses. Mechanical force is often referred as a regulating factor of musculoskeletal tissues (Pitsillides *et al*., 1999; Poulet *et al*., 2015; Sun, 2010; Zuscik *et al*., 2008). In addition, cartilage compression at physiological levels results in chondrocytes’ anabolic responses and are therefore considered protective (Zhang and Yao, 2021; Zuscik *et al*., 2008). It has been demonstrated that mechanical stimulation increases TGF-ß expression and signalling in cartilage and can be used to induce endochondral ossification processes (Zhang *et al*., 2015b; Zhen *et al*., 2021). But it is also clear that excessive or abnormal mechanical stimulation is detrimental. Indeed, these can induce catabolic processes in chondrocytes and are linked *in vivo* to OA development (Poulet *et al*., 2011; Qin *et al*., 2024; Zhang *et al*., 2022). Our data in TSK mice support these facts, linking excessive TGF-ß activity with increased mechanically-induced endochondral ossification. In addition, Fbn1 microfibrils harbour an RGD integrin-binding site suggesting a direct effect on mechano-signalling (Booms *et al*., 2005; Sakamoto *et al*., 1996; Zeyer *et al*., 2019); Fbn1 mutations in TSK mice may therefore result in abnormal mechanotransduction and increased joint pathology. Confirmation of the mechanisms by which FBN1 mutation in TSK mice are responsible for increased endochondral ossification in articular tissues in necessary.

In this study, we show that the effects of FBN1 mutation on articular soft tissue ossification is sex specific, affecting mostly males. Sexual dimorphism in FBN1 related pathologies have been reported before, in a tissue specific manner. While males reported to be more susceptible to the cardiovascular events in Marfan (Groth *et al*., 2017; Nucera *et al*., 2022), the skeletal manifestations show more severe symptoms in females (Taniguchi *et al*., 2023). A study by on a Danish cohort reported that risks for some musculoskeletal manifestations such as scoliosis and rheumatic diseases are significantly increased in women compared to men (Andersen *et al*., 2022). These sex differences may be attributed to a complex combination of several causes including genetics, environmental factors, and sex hormones. Estrogen is reported to have protective effects on vascular system (Mendelsohn, 2002) and may be a key player in reduced risk of cardiovascular diseases in women (Huang and Kaley, 2004), whilst high levels of testosterone are linked with cardiac remodelling and aortic enlargement (Cavasin *et al*., 2006). In addition, TGF-ß signalling pathway plays a crucial role in pathogenesis of Marfan Syndrome, which may be regulated differently in males and females.

Similarities between skeletal phenotypes are supported by mouse models with clear bone overgrowth and thinning in both sexes (Barisic-Dujmovic *et al*., 2007). In contrast, specific tissues show a sex effect including abnormal fat deposition, insulin resistance and metabolic adaptations (Muthu *et al*., 2022; Tiedemann *et al*., 2022). A few studies have mentioned the potential effects of estrogen on FBN1 expression and deposition (Natoli *et al*., 2005; Renard *et al*., 2017), which may contribute to some of these sex differences. FBN1 associated joint pathologies, such as those described in this study, however, have not yet been investigated and their potential sex differences unknown.

In parallel, joint pathology such as OA development do show a clear sex effect, mostly linked to menopause; indeed, women show higher incidence of OA after menopause (Laitner *et al*., 2021; Overstreet *et al*., 2023), whilst ovariectomised female mice showed increased cartilage damage (Gilmer *et al*., 2023; Sniekers *et al*., 2008). In contrast, young virgin mice show relative protection to OA development compared to their male counterparts, suggesting a potential protective effect of female hormones in joint pathologies. Unfortunately we were not able to assess OA development in our female TSK mice due to early elimination of our colonies due to COVID restrictions. The relative protection female TSK mice compared to male TSK on articular soft tissue ossification may include the interaction between TGF-ß and estrogen, with some suggestions of an inhibitory effect of estrogen on TGF-ß (Mao *et al*., 2019; Nasatzky *et al*., 1999; Yang *et al*., 2022). A study on TSK mice by Avouac et al. (Avouac *et al*., 2020) presented that estrogens are inhibiting TGF-ß induced dermal fibrosis through estrogen receptor alpha. This could be one of the factors that females are being protected from the actions of overactive TGF-ß in our TSK female mice. Further work is needed to better understand the effects of sex and osteoarthritis development, and the cellular mechanisms involved.

In summary, we describe for the first time a sex-specific effect of FBN1 mutation in TSK mice on joint pathology, especially articular soft tissue ossification, that may be linked to increased active TGF-ß and their interplay with estrogen inhibitory effects.

## Acknowledgements

We are grateful to Arthritis Research UK (20859), Versus Arthritis (22451), Chinese Scholarship Council, the Medical Research Council (MR/X021068/1), The MRC-Arthritis Research UK Centre for Integrated Research into Musculoskeletal Ageing (CIMA), and the Institute of Aging and Chronic Disease (University of Liverpool) for providing funding for this study.

## Authors contributions

B.P., D.A. and G.B-G contributed to the design of this work. C.K., X.W., T.D., Y.W., L.R-M and E.S. contributed to the experimental work, data collection and contributed to analysis and interpretation of data. B.P., C.K. and T.D. drafted the work. X.W., Y.W., L.R-M, E.S., D.A. and G.B-G revised critically for important intellectual content. All authors read and approved the final manuscript. All authors agree to be accountable for all aspects of the work in ensuring that questions related to the accuracy or integrity of any part of the work are appropriately investigated and resolved.

